## Supplemental Figures and Materials for "An abundant quiescent stem cell population in *Drosophila* Malpighian tubules protects principal cells from kidney stones"

#### SUPPLEMENTAL ONLINE MATERIAL

##### Supplemental Figures

**Figure S1. Adult *Drosophila* Malpighian tubules are organized into different compartments, Related to Figure 1**

(A) Schematic drawing of adult Malpighian tubules showing the compartments and some useful Gal4 drivers. (B) *31E09-Gal4> GFP* marks the lower ureter. (C) *Pvr-GFP* is expressed in PCs of lower ureter. (D) *c507 >RFP* marks the ureter as well as lower tubules. (E) *Uro>RFP* marks main segment. (F) Stellate cells marked by the expression of *tsh-Gal>RFP* are only present in the upper tubules. *Drosophila* ureter is surrounded by circular muscle and longitudinal muscle as revealed by phalloidin staining (green).

**Figure S2. No replenishment of either stellate cells or principal cells in the upper tubules after ablation, Related to Figure 3**

(A-B) Expression of *rpr* and *hid* with *c507-Gal4*$^{ts}$ led to SCZ regeneration. (A) Control. (B) Principal cells in the SCZ were ablated by the expression of *rpr* and *hid* at 29$^{o}$C for 7 days. After recovery at 18$^{o}$C for 21 days, all big preexisting principal cells (marked by high expression levels of Cut, in red) were gone and replaced by smaller principal cells.

(C) Both *tsh-Gal4>RFP* (red) and *tsh-lacZ* (green) express in stellate cells.

(D-G) Stellate cell ablation and recovery. Stellate cells were readily present in control animals (D) as revealed by the expression of *tsh-lacZ* with X-gal staining. After expression of *UAS-hid,rpr* driven by *tsh-Gal4*$^{ts}$ for 7 days at 29$^{o}$C (E), most stellate cells were ablated. (F) No replenishment of stellate cells was detected at day 21 after shifting stellate-cell ablated animals back to 18$^{o}$C for recovery. (G) Quantification of the stellate cell number in each upper tubule. Data represent means± SD, n=20-30 animals.

(H-I) Main segment principal cell ablation and recovery. (H) *Uro-Gal4* driven *UAS-RFP* was specifically expressed in the principal cells at the main segment. Inset : saggittal section view showing *Uro-Gal4* driven *UAS-RFP* was not expressed in stellate cells (arrowhead). Scale bar in inset: 20 um. (I) Adults with the genotype of *UAS-rpr,UAS-hid/+; tub-Gal80*$^{ts}$*/Uro-Gal4,UAS-myrRFP* were shifted to 29$^{o}$C for 7 days and then shifted back to 18$^{o}$C for recovery for 21 days. Increased cell density was observed near the boundary (indicated by yellow arrowheads) between lower tubule and main segment, whereas drastically reduced cell density was observed in the distal tubule (indicated by white triangles).

**Figure S3. Damage activates multiple proliferation pathways, Related to Figure 3**

(A-D) Surgical resection of one Malpighian tubule was used to induce damage prior to assaying various signaling pathway activity. The intact Malpighian tubule was used to serve as an internal control. (A) Expression of *puc*$^{E69}$*-Gal4>UAS-GFP* was induced in cells near the surgical site 24 hours after surgery. (B) Expression of *10XStat-GFP* was upregulated in cells near the surgical site 2 days after surgery. (C) dpErK expression was specifically upregulated in RSCs near the surgical site 2 days after surgery. (D) *Diap1-lacZ* was upregulated near the surgical site 4 days after surgery.

**Figure S4. Forced expression of NICD or *ct* is sufficient to drive RSC differentiation, Related to Figure 4**

(A-C) *esg-Gal4*$^{ts}$ directed Flp-out lacZ marked lineage tracing in control (A), and following expression of NICD (B) or *ct* (C) for 14 days. RFP marks *esg*$^{+}$ cells, lacZ marks all the cells derived from *esg*$^{+}$ cells. (D-E) X-gal staining showing expression of *Alp4* in control Malpighian tubules (D), and following expression of NICD (E) or *ct* (F) driven by *esg-Gal4*$^{ts}$ for 7 days. Note the small ectopic *Alp4*$^{+}$ cells (denoted by yellow arrowheads) in (E) and (F).

**Figure S5. esg+ cells are largely arrested at G0 and G2 phases in normal condition. (**A) The Fly-FUCCI system consists of fluorescently tagged E2F1 and CycB degrons. Predicted expression of Fly-FUCCI in different cell cycle phases is shown. (B) Expression of Fly-FUCCI (CFP-E2F1$_{1\text{-}230}$;Venus-CycB$_{1\text{-}266}$) driven by *esg-Gal4,UAS-myrRFP*. The red triangles denote the G0 *esg>RFP+* cells that are double-negative for CFP-E2F1$_{1\text{-}230}$ and Venus-CycB$_{1\text{-}266}$ , and the yellow arrowheads denote the G2 *esg>RFP+* cells that are double-positive for CFP-E2F1$_{1\text{-}230}$ and Venus-CycB$_{1\text{-}266}$. Cyan arrows denote G1 *esg>RFP+* cells that are positive for CFP-E2F1$_{1\text{-}230}$ and negative for Venus-CycB$_{1\text{-}266.}$ Green arrow denotes an S phase *esg>RFP+* cells that is positive for Venus-CycB$_{1\text{-}266}$ and negative for CFP-E2F1$_{1\text{-}230.}$ (C) Enlarged view of boxed regions in (B). Note the G0 cells have a 2C DNA content while G2 cells have a 4C DNA content.

**Figure S6. RSCs respond to stone formation**

(A) *Drosophila ry* gene encodes xanthine dehydrogenase (XDH), which catalyzes the conversion of hypoxanthine and xanthine to uric acid. Allopurinol is an inhibitor of XDH. (B-D) Stones progressively developed in *ry*$^{506}$ flies and in flies reared on allopurinol food. (B) Representative DIC images of *ry*$^{506}$ ureter and lower tubules. Note the stones do not transmit light and hence appear dark. (C and D) Allopurinol feeding recapitulated stone development caused by loss of *ry*. (E) Immunofluorescence micrograph showing a 7 day-old *ry*$^{506}$ fly carrying stones (indicated by yellow arrowheads) in the lower tubules and ureters. Note that the SCZ bearing larger stones and more stones contains significantly more cells. (F) Scatter plot showing the correlation between stone sizes (area) and the number of cells present in the SCZ from 7 day-old $ry^{506}$ flies. (G-H) EdU incorporation in $esg^+$ cells from *esg>RFP* flies fed on control diet (G) and allopurinol augmented food (H). Note significantly more $esg^+$ cells are $EdU^+$ in allopurinol-treated flies. (I) Fly-FUCCI system was used to assess cell cycle phase distribution of $esg^+$ cells after allopurinol treatment. (J-K) MARCM clones showing clones were barely detectable in control animals with a low dose of heat shock (J), Allopurinol administration increased clone induction in the SCZ with a low dose of heat shock (K). (L) Quantification of MARCM clone numbers in the SCZ with 2X heat shocks. Data are means± SD, *** denotes Student's t test $p<0.001$. n=20-49. (M) Distribution of marked clones in Malpighian tubule 14 days after clone induction.

**Figure S7. Stone-induced damage causes activation of multiple proliferation pathways**

(A) Volcano plot showing differentially expressed genes between *ry* mutant lower tubules carrying stones and wild-type lower tubules. Genes with fold change more than 2 and Padj < 0.05 are in pink. The table on the right shows the pathways that the highlighted genes are involved in the volcano plot. (B) Expression of $puc^{E69}$*-Gal4 >GFP* in one pair of Malpighian tubules carrying stones (denoted by yellow arrowhead) and the other pair of Malpighian tubules without stones (red triangle) from the same animal after allopurinol treatment. (C) Expression of Jak/Stat pathway reporter, *10XStat92E-GFP* in control Malpighian tubules (left panels) and in Malpighian tubules carrying stones induced by allopurinol (right panels). (D-E) X-gal staining showing the expression of *Diap1-lacZ* (D) and *ex-lacZ* (E) in normal and stone-carrying Malpighian tubules. (F-G) Expression of Notch signaling reporter, *NRE-GFP* in $ry^{+/-}$ Malpighian tubules (F) and in $ry^{506}$ Malpighian tubules bearing stones.

### STAR★METHODS

#### KEY RESOURCES TABLE

| REAGENT or RESOURCE | SOURCE | IDENTIFIER |
|---|---|---|
| Antibodies | | |
| Mouse anti-Cut | Developemental Studies Hybridoma Bank (DSHB) | Cat#2B10; RRID: AB_528186 |
| Mouse anti-Delta | DSHB | Cat#C594.9B; RRID: AB_528194 |
| Mouse anti-NECD | DSHB | Cat#C458.2H; RRID: AB_528408 |
| Rabbit anti-β-Gal | Cappel | Cat#55976; AB_2313707 |
| Mouse anti-β-Gal | Promega | Cat#Z3781; AB_430877 |
| Rabbit anti-GFP | ThermoFisher | Cat# A-11122; RRID: AB_221569 |
| Mouse anti-PH3 | Cell Signaling Technology | Cat# 9706, RRID:AB_331748 |
| Mouse anti-dpErK | Sigma-Aldrich | Cat# M9692, RRID:AB_260729 |
| Rabbit anti-RFP | Rockland | Cat# 600401379, RRID: AB_2209751 |
| Mouse anti-rat-CD2 | Bio-Rad | Cat# MCA154R, RRID: AB_321239 |
| Alexa 488 goat anti-Mouse | ThermoFisher | Cat#: A11001; RRID: AB_2534069 |
| Alexa 568 goat anti-Mouse | ThermoFisher | Cat#: A11004; RRID: AB_ 2534072 |
| Alexa 488 goat anti-Rabbit | ThermoFisher | Cat#: A11034; RRID: AB_2576217 |
| Alexa 568 goat anti-Rabbit | ThermoFisher | Cat#: A11041; RRID: AB_ 2534098 |
| Experimental Models: Organisms/Strains | | |
| *D.melanogaster*: $N^{55e11}$ $FRT^{19A}$/*FM7c* | Bloomington Drosophila Stock Center (BDSC) | RRID: BDSC_28813 |
| *D.melanogaster*: $y^{1}$;; $ry^{506}$ | BDSC | RRID: BDSC_4405 |
| *D.melanogaster*: Oregon-R | BDSC | RRID: BDSC_25211 |
| *D.melanogaster*: NRE-EGFP | BDSC | RRID: BDSC_30727 |
| *D.melanogaster*: UAS-N.intra | BDSC | RRID: BDSC_52008 |
| *D.melanogaster*: 10X Stat92E-GFP | BDSC | RRID: BDSC_26198 |
| *D.melanogaster*: tsh-lacZ | BDSC | RRID: BDSC_11370 |
| *D.melanogaster*: Alp4-lacZ | BDSC | RRID: BDSC_12285 |
| *D.melanogaster*: Diap1-lacZ | BDSC | RRID: BDSC_12093 |
| *D.melanogaster*: ex-lacZ | BDSC | RRID: BDSC_44248 |
| *D.melanogaster*:UAS-myr::tdTomato | BDSC | RRID: BDSC_32222 |
| *D.melanogaster*: UAS-ct | BDSC | RRID: BDSC_36496 |
| *D.melanogaster*: UAS-rpr,UAS-hid | J. Nambu | (Zhou et al., 1997) |
| *D.melanogaster*: UAS- $N^{RNAi}$ | BDSC | RRID: BDSC_35640 |
| *D.melanogaster*: UAS- Rac1.N17 | BDSC | RRID: BDSC_6292 |
| *D.melanogaster*: $FRT^{19A}$ | BDSC | RRID: BDSC_1709 |
| *D.melanogaster*: $FRT^{42D}$ | BDSC | RRID: BDSC_1802 |
| *D.melanogaster*: Act5C>stop>lacZ | BDSC | RRID: BDSC_6355 |
| *D.melanogaster*: UAS-Flp | BDSC | RRID: BDSC_4539 |
| *D.melanogaster*: $Puc^{E69}$-Gal4 | BDSC | RRID: BDSC_6762 |
| *D.melanogaster*: c507-Gal4 | BDSC | RRID: BDSC_30840 |
| *D.melanogaster*: Uro-Gal4 | BDSC | RRID: BDSC_44416 |
| *D.melanogaster*: tsh-Gal4 | BDSC | RRID: BDSC_3040 |
| *D.melanogaster*: P{tubP-GAL80[ts]}20 | BDSC | RRID: BDSC_7019 |
| *D.melanogaster*: esg-Gal4 | N. Perrimon | (Micchelli and Perrimon, 2006) |
| *D.melanogaster*: UAS-myr::tdTomato | BDSC | RRID: BDSC_32222 |
| *D.melanogaster*: P{tubP- | BDSC | RRID: BDSC_7018 |

| | | |
|---|---|---|
| GAL80[ts]}ncd[GAL80ts-7] | | |
| *D.melanogaster:* cad-Gal4 | BDSC | RRID: BDSC_3042 |
| *D.melanogaster:* UAS-rCD2.RFP,UAS-GFPi, $FRT^{40A}$ | BDSC | RRID: BDSC_56184 |
| *D.melanogaster:* UAS-rCD8.GFP,UAS-rCD2i,$FRT^{40A}$ | BDSC | RRID: BDSC_56185 |
| *D.melanogaster:* UAS-CFP.E2f1.1-230; UAS-Venus.CycB.1-266 | BDSC | RRID: BDSC_55102 |
| *D.melanogaster:* Df31-GFP | Lab stock | (Buszczak et al., 2007) |
| *D.melanogaster:* Pvr-sGFP | Vienna *Drosophila* Resource Center | RRID: VDRC_318162 |
| *D.melanogaster: UAS-cd8GFP hsFlp; $FRT^{42D}$ tub-Gal80; tub-Gal4/TM6B, Tb* | Lab stock | *N/A* |
| *D.melanogaster:* UAS-upd1 | J. Urban (X. Chen lab, JHU) | N/A |
| Chemicals, Peptides, and Recombinant Proteins | | |
| DAPI | Sigma-Aldrich | Cat#D9542 |
| Allopurinol | Sigma-Aldrich | Cat#A8003 |
| collagenase | Sigma-Aldrich | Cat#C2674 |
| elastase | Sigma-Aldrich | Cat#E0258 |
| FBS | ThermoFisher | Cat#16140063 |
| TRIzol | Invitrogen | Cat#15596026 |
| TruSeq RNA Library Prep Kit v2 | illumina | RS-122-2001 |
| Critical Commercial Assays | | |
| Click-iT™ EdU Cell Proliferation Kit for Imaging, Alexa Fluor™ 488 dye | ThermoFisher | Cat#C10337 |
| Software and Algorithms | | |
| Adobe Illustrator CC 2018 | Adobe | https://www.adobe.com/products/illustrator.html |
| IMARIS v9.2.1 | Bitplane | https://imaris.oxinst.com |
| Fiji | NIH | https://fiji.sc/ |
| HISAT2 v2.1.0 | (Pertea et al., 2016) | https://ccb.jhu.edu/software/hisat2/index.shtml |
| DESeq2 | (Love et al., 2014) | https://bioconductor.org/packages/release/bioc/html/DESeq2.html |
| CellRanger v2.1.1 | 10x Genomics Inc. | https://support.10xgenomics.com/single-cell-gene-expression/software/pipelines/latest/what-is-cell-ranger |
| Seurat v2.3.4 | (Butler et al., 2018) | http://satijalab.org/seurat/ |
| R v3.5.2 | R core team | https://www.r-project.org |

#### CONTACT FOR REAGENT AND RESOURCE SHARING

#### EXPERIMENTAL MODEL AND SUBJECT DETAILS

##### *Drosophila* and Husbandry

The *Drosophila* stocks used are listed in the resource table. Flies were reared on normal cornmeal molasses food and maintained at room temperature( 23-25°C)  unless otherwise specified.

#### METHOD DETAILS

##### Immunostaining and microscopy

Malpighian tubules were dissected in Grace's insect buffer and left attached to *Drosophila* gut. Then fixed in 4% paraformaldehyde on a nutator at room temperature for 20min. Sample were then washed 3 times in PBT (1XPBS + 0.1%TritonX-100) and blocked in PBT plus 5% normal goat serum (NGS) for 1h followed by primary antibodies incubation overnight at 4$^o$C. Secondary antibodies were incubated for 2-3 hours at room temperature or overnight at 4$^o$C. In order to minimize the autofluorescence elicited by ureter stones, an alternative staining procedure was used. Malpighian tubules bearing ureter stones were dissected and fixed in 800ul fixative solution (containing 100ul 16% EM-grade paraformaldehyde + 300ul Grace's insect buffer + 400ul n-heptane) on a nutator at room temperature for 20min.  After fixation, the aqueous phase was removed and 400 ul 100% methanol was added, followed by vigorous hand shaking for 30s. Samples were subsequently washed with 100% and 50% methanol for 5 min respectively. Samples were then washed 3 times in PBT (1XPBS + 0.1%TritonX-100) and blocked in PBT plus 5% NGS for 1h followed by primary antibodies overnight at 4$^{o}$C. Secondary antibodies were incubated for 2-3 hours at room temperature or overnight at 4$^{o}$C. Samples were washed 3 times in PBT, 5 minutes each after antibody incubation. Finally, DNA were stained with DAPI (100ng/ml) in PBT for 5min. Samples were mounted in 50% glycerol and imaged with a Leica TCS SP5 or Leica TCS SP8 confocal microscope.

**Electron microscopy**

Malpighian tubules were dissected in Grace's Buffer and the gut and upper tubules were carefully removed. Samples were fixed for 1 hour in fixative solution (3% glutaradehyde + 1% formaldehyde +0.1 M cacodylate buffer + Ca+Mg, pH=7.4). Following rinsing with 0.1 M cacodylate buffer, samples were embedded in agarose at 50$^{o}$C, washed with cacodylate buffer three times for10 minutes each time, then the samples were further fixed in 1% $OsO_4$ + 1% KFeCN in cacodylate buffer for 45 min. After rinsing in $H_2O$ twice for 10 minutes each time, samples were rinsed in 0.05 M maleate (pH 6.5) for 10 min and stained for 1.5 hr in 0.5% uranyl acetate, 0.05 M maleate (pH 6.5). After rinsing in $H_2O$ for 10 minutes, samples were dehydrated in an ethanol series (35% twice for 10 minutes, 50% for 10 minutes, 75% for 10 minutes, 95% for 10 minutes and 100% three times for 15 minutes), followed by incubating in propylene oxide 4X 15 minutes each, and then in 1:1 propylene oxide : epoxy resin for 1 hr. After three changes of 100% resin (1 hour each), resin was allowed to harden overnight at 55$^{o}$C and then at overnight 70$^{o}$C. Electron microscopy images were captured with a Phillips Tecnai 12 microscope and recorded with a GATAN multiscan CCD camera using Digital Micrograph software.

**Heat shock scheme for mosaic analysis**

MARCM systems were used to generate mitotic clones (Lee and Luo, 1999). Flies with appropriate genotypes were put in empty vials augmented with yeast paste on the wall and heat-shocked in a 37$^{o}$C water bath once to multiple times (60 min each time), with an interval of 24 hours between heat shocks. To assay the proliferative activity of RSCs upon injury caused by ureter stones, newly eclosed female flies were reared on cornmeal-molasses food containing 3mM Allopurinol for 7 days prior to heat shock. After 2X heat shocks, flies were transferred to Allopurinol food and transferred every 2-3 days.

For twin-spot MARCM (Yu et al., 2009), animals were subjected to surgical injury. One of the anterior pair of Malpighian tubules was severed in the SCZ. After recovery for 1 day, animals were heat-shocked in a 37$^{o}$C water bath for 60 min. The resultant RFP-marked and GFP-marked twin-spots were scored as "symmetric" if both clones contained multiple cells at day 5-7 after clone induction. Twin spots were scored as "asymmetric" if they consisted of one polyploid cell in one color and multiple cells in the other color.

**Surgical removal of *Drosophila* Malpighian Tubule**

Flies were anesthetized on a $CO_2$ pad prior to surgery, and were laid on their sides to expose the boundary between the dorsal and ventral abdomen. The A1 abdominal pleura was opened up with fine forceps (Fine Science Tools #5SF) under a stereoscopic microscope and one of the anterior pair of Malpighian tubules was pulled out and severed at appropriate sites as indicated in the figure legends. After surgery, flies were transferred to regular food for recovery.

**Flip-out lacZ marked lineage tracing**

To carry out Flip-out lacZ marked lineage tracing of RSCs after surgical injury, *Act5C>STOP>lacZ* tracer flies were crossed to *esg-Gal4,UAS-flp,UAS-myrRFP; tub-Gal80$^{ts}$* driver flies at 18$^{o}$C. 3-7 day old progeny flies with appropriate genotypes were subject to surgical removal of one of the Malpighian tubules at desired sites, as specified in the figure legends prior to being shifted to 29°C to initiate lineage labeling. In order to conduct Flip-out lacZ marked lineage tracing of RSCs expressing UAS-$N^{RNAi}$, UAS-NICD or UAS-ct, *Act5C>STOP>lacZ* was combined with the transgene of interest first. The resultant *UAS-X; Act5C>STOP>lacZ* tracer flies were then crossed to the *esg-Gal4,UAS-flp,UAS-myrRFP; tub-Gal80$^{ts}$* driver flies.

**Allopurinol feeding**

Allopurinol stock solution (300 mM) was made by dissolving 0.41g of Allopurinol (Sigma-Aldrich, cat# A8003-25g) in 10 ml of 1M sodium hydroxide. Normal cornmeal molasses food was melted and supplemented with 1/100 volume of allopurinol stock solution. After well mixed, the allopurinol food was poured into individual vials. Female flies with appropriate genotypes were cultured on allopurinol food as indicated in figure legends.

**Single cell RNA-seq**

*Drosophila* Malpighian tubules from 5-7 day old Oregon-R female flies were dissected in nuclease-free PBS on ice. *Drosophila* gut and upper tubules were carefully removed. Ureter and lower tubules from ~200 females were collected within 2 hours and then transferred to an Eppendorf tube containing 500 μl of dissociation buffer (1 mg/ml collagenase, 0.5 mg/ml elastase in nuclease free PBS). After incubating at 27°C for 1 hour on a nutator, the digestion reaction was stopped using 100 μl of fetal bovine serum. The dissociated cell suspension was then passed through a 70 μm cell strainer. The cells were spun down at 500g, 4°C for 10 min and the supernatant was removed. 1 ml ice cold PBS was added to briefly resuspend and rinse cell pellets. After rinsing, the cells were spun down and resuspended with 100 μl of nuclease-free PBS.

Cells were encapsulated and the cDNA library was prepared at Genetic Resources Core Facility at the Johns Hopkins School of Medicine. The cDNA library was sequenced on an Illumina NextSeq 500 sequencer at the Carnegie Institution. A total of ~126 M reads were acquired. Reads were mapped to the *Drosophila* melanogaster genome BDGP6 using CellRanger (10x Genomics Inc. version 2.1.1). Median UMI counts per cell was 8,737. Mean reads per cell was 173,493 and the median genes per cell was 1,662. R package Seurat (2.3.4) was used to analyze single cell RNA results (Butler et al., 2018).

**Bulk RNA-seq**

*Drosophila* ureter and lower tubules were dissected from ~100 females in nuclease-free PBS on ice. Gut and upper tubules were carefully removed. The isolated tissues were transferred to 500 μl of ice cold TRIzol and frozen at -80$^{o}$C. When all replicates for ctrl and experimental groups were ready, the RNAs were extracted following manufacturer's instructions. cDNA libraries were prepared from poly(A)-selected RNA using Illumina TruSeq RNA Library Prep Kit v2, and sequenced using an Illumina Nextseq 500 sequencer. Single-end reads were mapped to the *Drosophila* melanogaster genome (dm6) using HISAT2 2.1.0 (Pertea et al., 2016) , transcripts were assembled using StringTie. Reads aligned to genes were calculated using featureCount (Liao et al., 2014) and differentially expressed genes were identified using R package DESeq2 (Love et al., 2014).

**Genetic cell ablation and recovery**

Cell-type specific genetic ablation was achieved by crossing *UAS-rpr,UAS-hid;tub-Gal80*$^{ts}$ with cell-type specific Gal4 lines. For ablation of principal cells in the SCZ, *c507-Gal4,UAS-*

*myr::tdTomato* males were crossed with *UAS-rpr,UAS-hid;tub-Gal80$^{ts}$* females at 18$^o$C. 3-5 day old *UAS-rpr,UAS-hid/+; tub-Gal80$^{ts}$/+; c507-Gal4,UAS-myr::tdTomato/+* females were then shifted to 29$^o$C for 7 days. After that, the flies were shifted back to 18$^o$C to minimize the expression of *rpr* and *hid* to recover. Similarly, *tsh-Gal4* and *Uro-Gal4* lines were used to ablate stellate cells and principal cells at the main segment of Malpighian tubules, respectively.

**X-Gal Staining**

Malpighian tubules were dissected into ice cold PBS and fixed in 0.5% Glutaraldehyde/PBS at room temperature for 15 min. After briefly wash in PBS, the samples were transferred into 1 ml of staining solution (10 mM $NaH_2PO_4 \cdot H_2O$, 10 mM $Na_2HPO_4 \cdot 2H_2O$, 150 mM NaCl, 1 mM $MgCl_2 \cdot 6H_2O$ , 3.1mM $K_4[Fe(CN)_6] \cdot 3H2O$, 3.1mM $K_3[Fe(CN)_6]$, 0.3% TritonX-100, 0.2% X-Gal) and incubated overnight at room temperature.

**EdU incorporation**

Depending on the design of experiments, different EdU pulse-chase strategies were used to assay EdU incorporation. To assay EdU incorporation in flies under normal condition, 3-5 day old females (*esg-Gal4,UAS-myr::tdTomato*) were fed on standard cornmeal molasses food supplemented with 0.5 mM EdU (Click-iT™ EdU Alexa Fluor™ 488 Imaging Kit, ThermoFisher) for 2-4 days. EdU incorporation were detected on day 2 and day 4 following manufacturer's instructions. To assay EdU incorporation in flies after surgical removal of upper Malpighian tubules, 3-5 day old females (*esg-Gal4,UAS-myr::tdTomato*) were first fed on EdU food for 2 days and were continuously cultured on EdU food for 2 more days after amputation of Malpighian tubules.

#### QUANTIFICATION AND STATISTICAL ANALYSIS

##### Nuclear volume measurement

z-stack images were acquired using a Leica TCS SP8 confocal microscope with a 20X objective lens (NA=0.75). 3D images were reconstructed using Bitplane IMARIS v9.2.1 (http://www.bitplane.com/). Surface reconstruction of the DNA (DAPI staining) was created using surface tool of IMARIS. Nuclear volume was calculated by IMARIS in the surface statistics.

##### Lifespan measurement

Wandering L3 larvae of control (*esg*$^{ts}$ *>RFP*) and *esg*$^{ts}$ *>RFP + Rac1.N17* were sorted and shifted from 18$^{o}$C to 29$^{o}$C. After eclosion, groups of 15-20 females of each genotype were placed into separate vials with normal cornmeal food or Allopurinol-augmented food and maintained at 29$^{o}$C. The flies were transferred to new food every other day. The number of dead flies were recorded and the number of flies that escaped during transfer was excluded from the study. The statistical significance was determined by log-rank tests. ns denotes $p > 0.05$, * denotes $p < 0.05$, ** denotes $p < 0.01$, *** denotes $p < 0.001$.

Figure S1 related to Figure 1

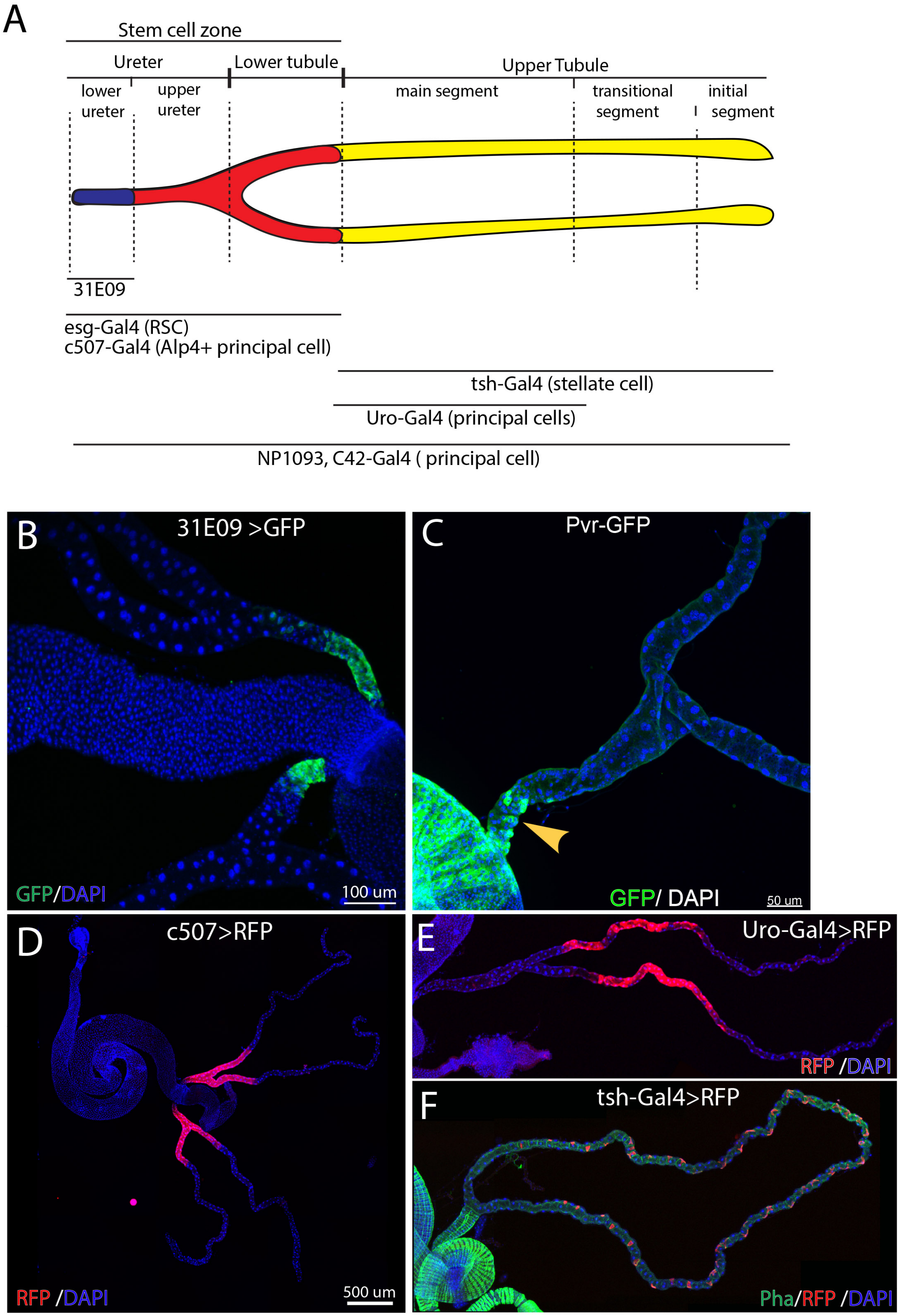

Figure S2 related to Figure 3

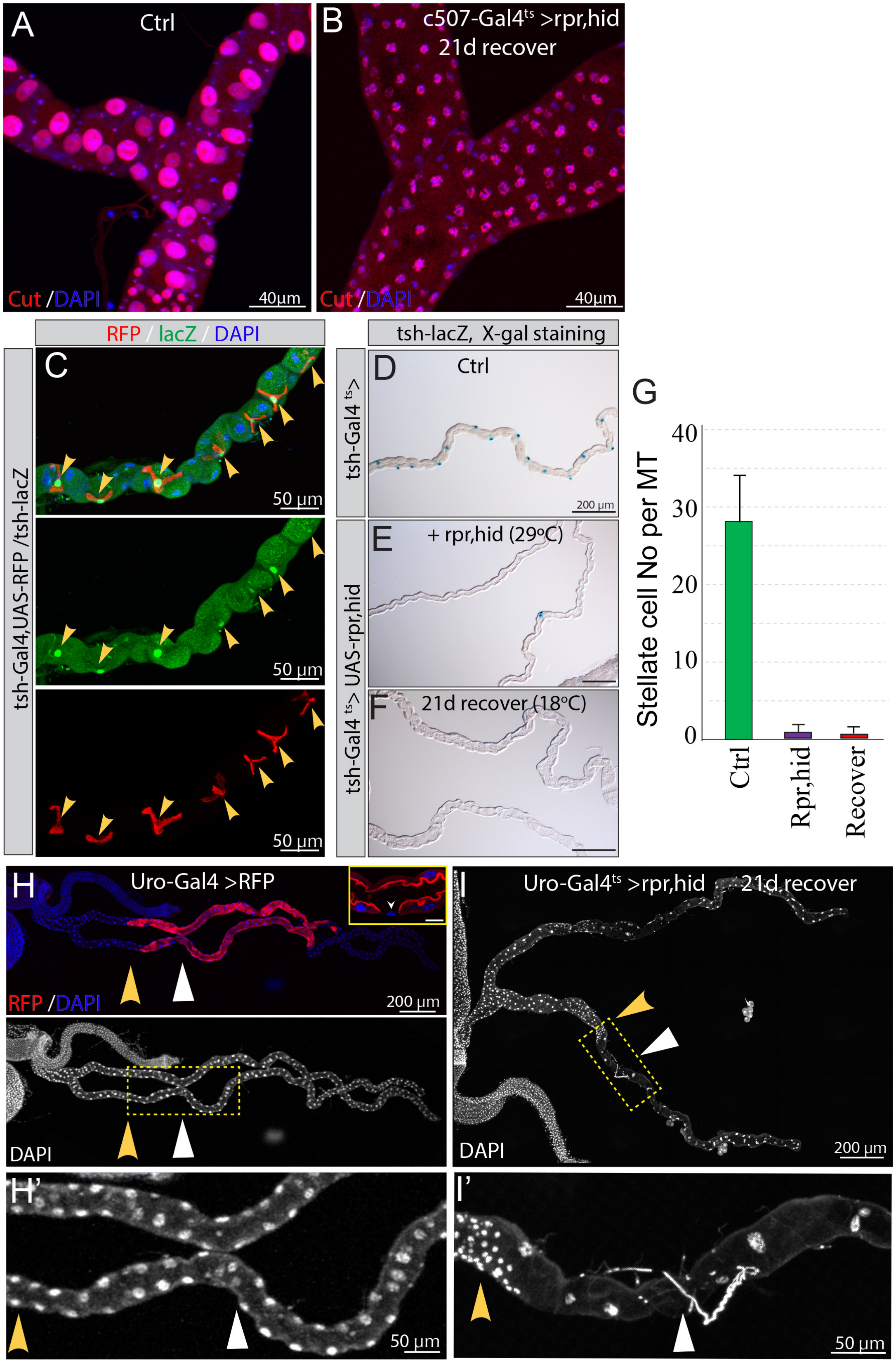

Figure S3 related to Figure 3

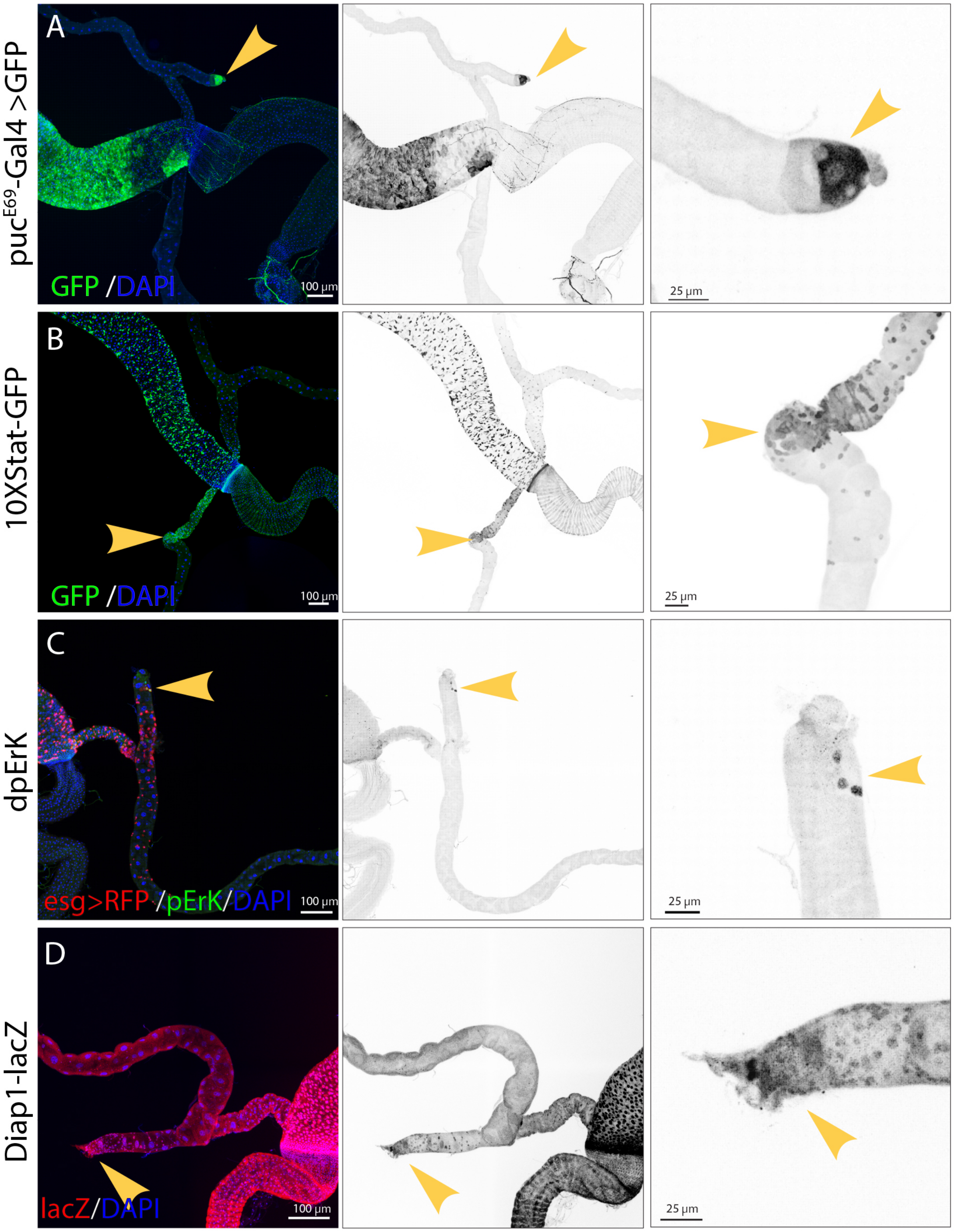

Figure S4 related to Figure 4

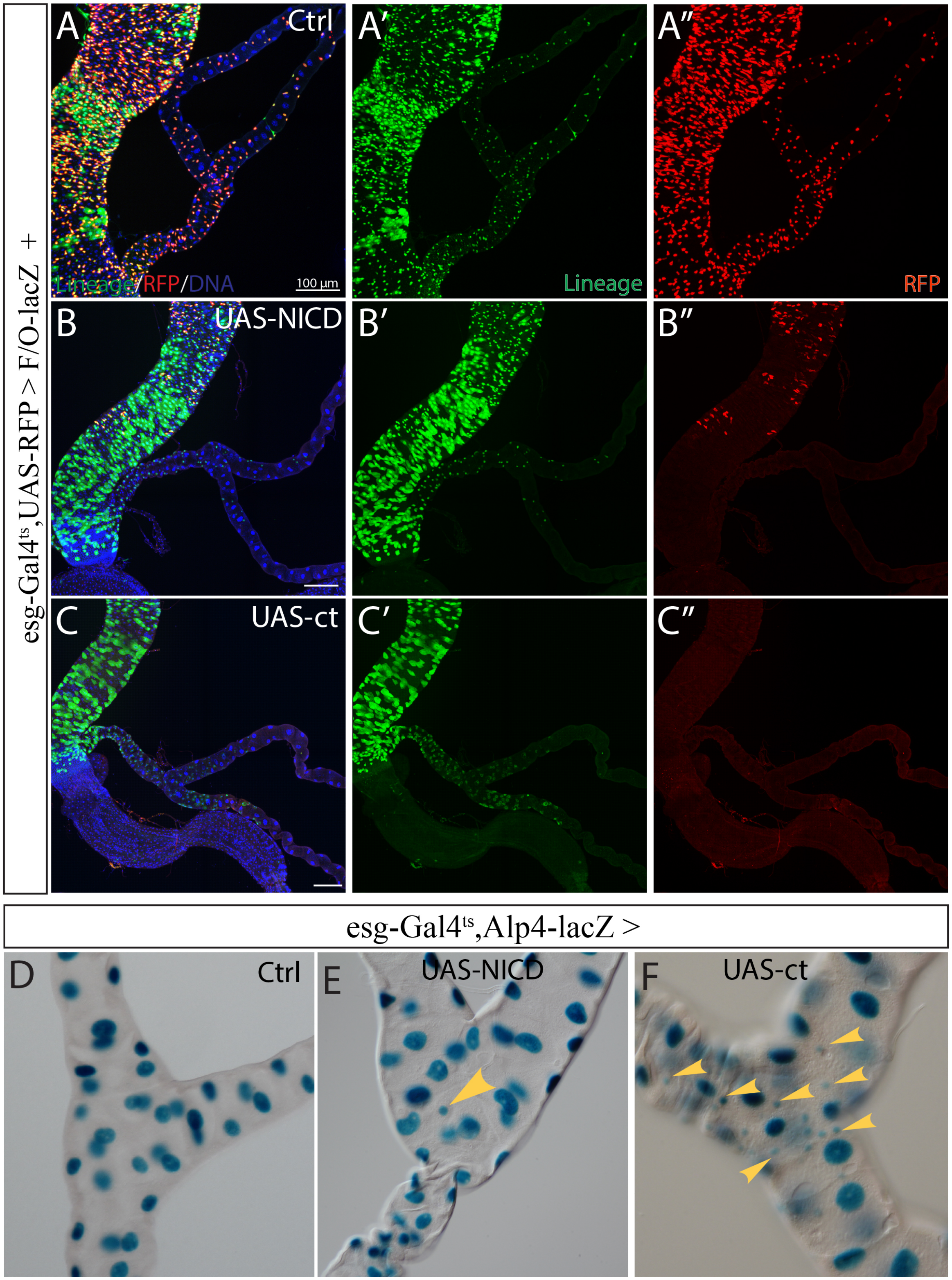

Figure S5 related to Figure 6

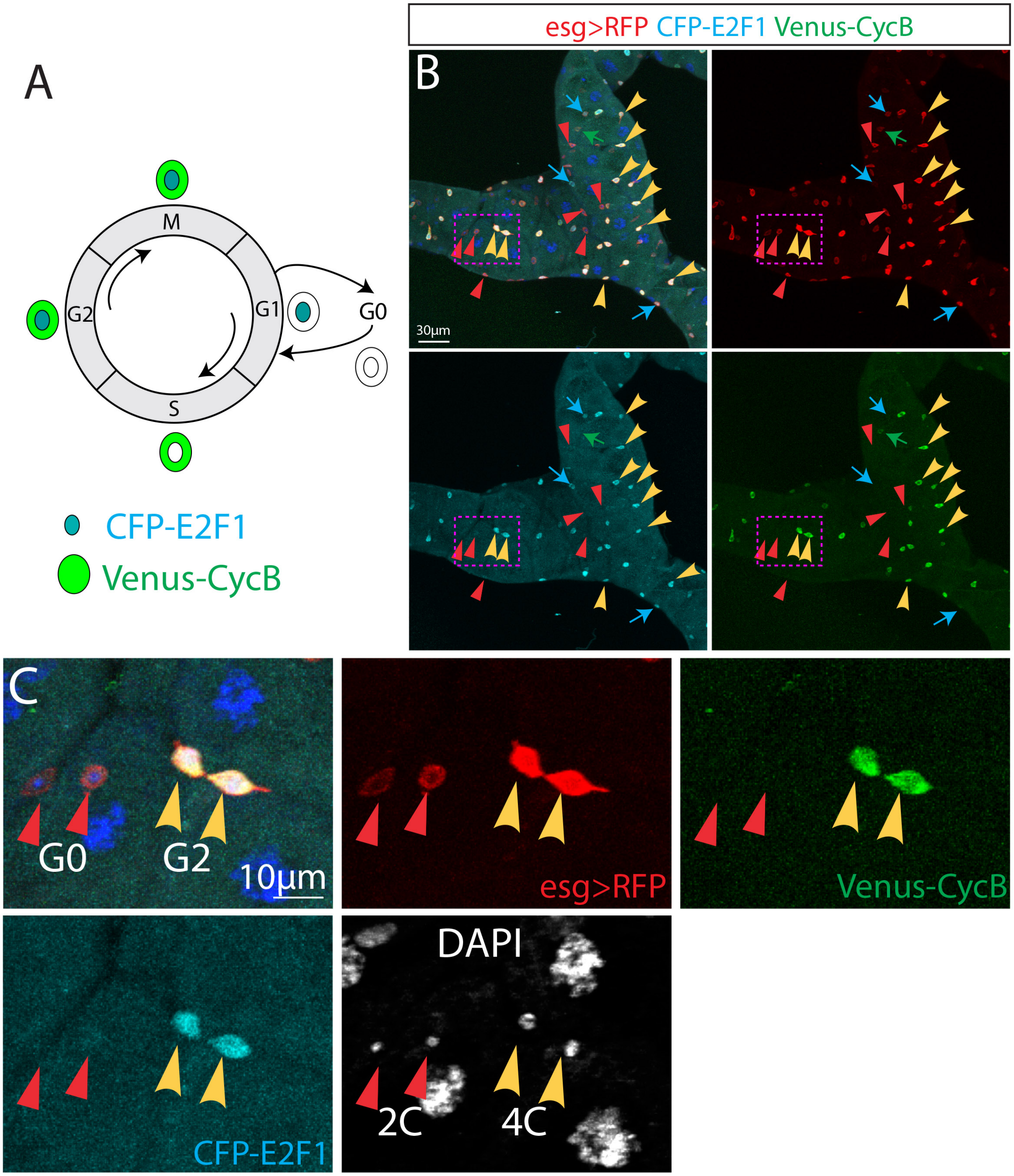

Figure S6 related to Figure 7

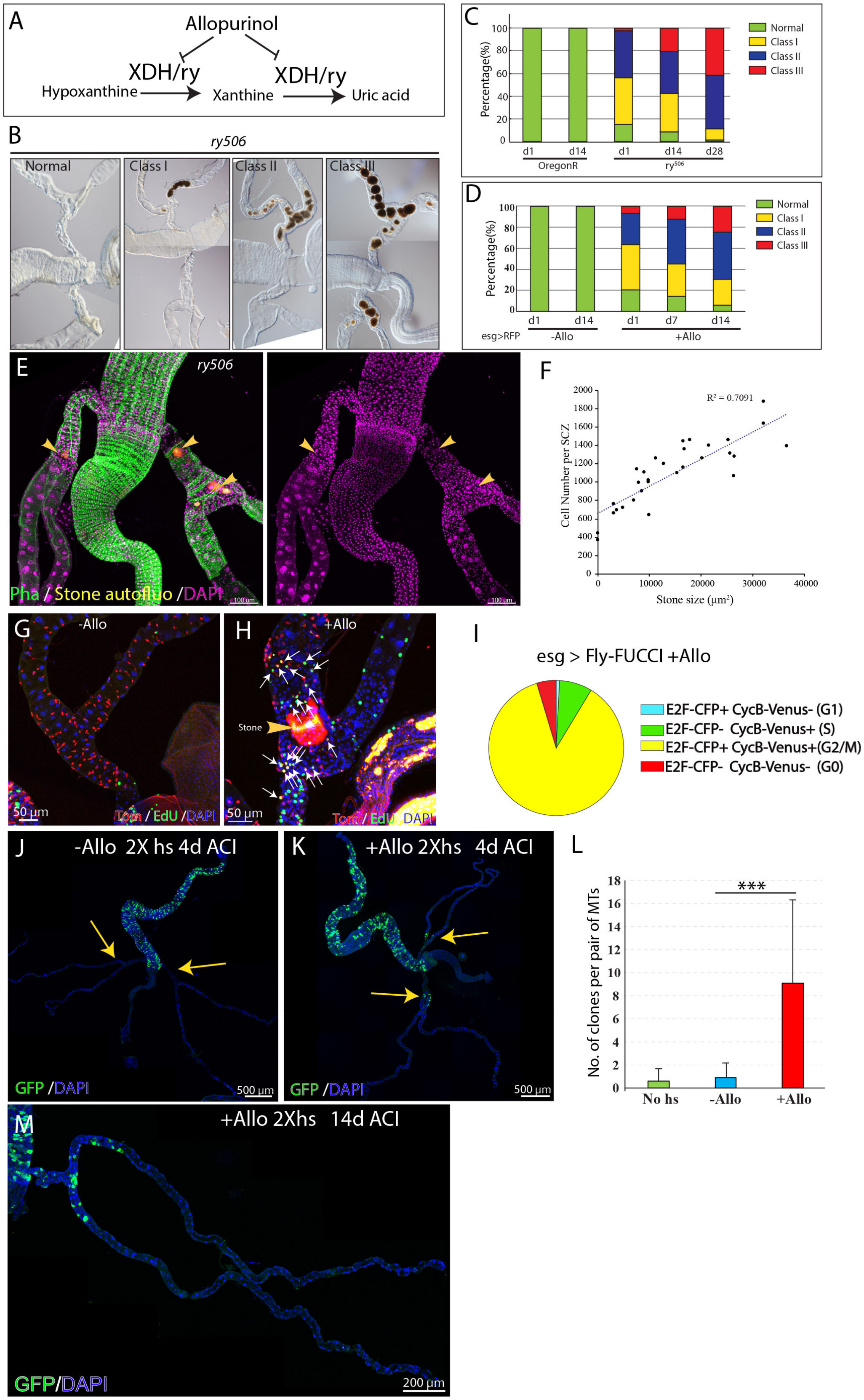

Figure S7 related to Figure 7

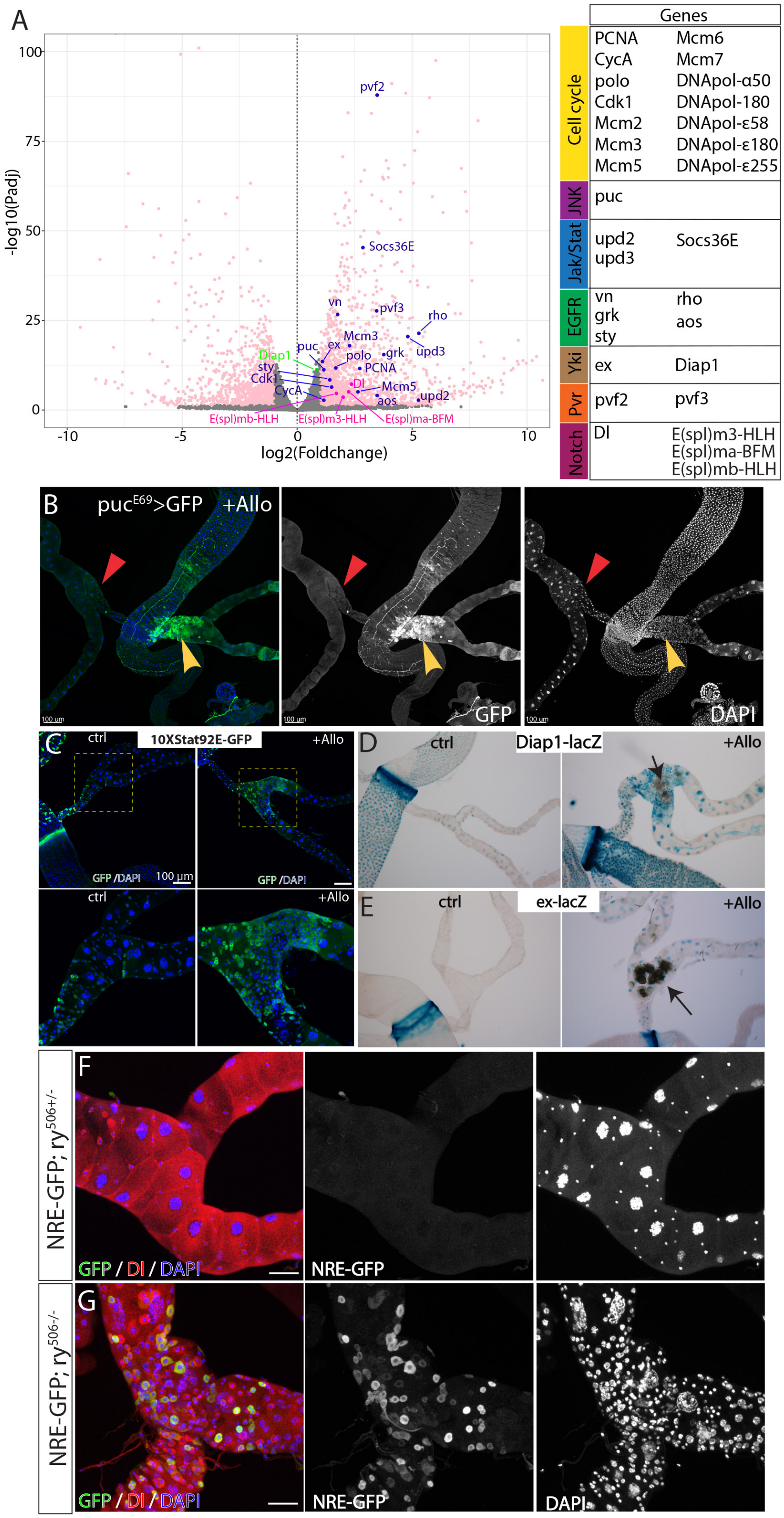
